# SPIN - Species by Proteome INvestigation

**DOI:** 10.1101/2021.02.23.432520

**Authors:** P.L. Rüther, I.M. Husic, P. Bangsgaard, K. Murphy Gregersen, P. Pantmann, M. Carvalho, R.M. Godinho, L. Friedl, J. Cascalheira, M.L.S. Jørkov, M.M.. Benedetti, J. Haws, N. Bicho, F. Welker, E. Cappellini, J.V. Olsen

## Abstract

Genetic species determination has become an indispensable tool in forensics, archaeology, ecology, and food authentication. The available methods are either suited for detecting a single taxon across many samples or for screening a wide range of species across a few samples. Here, we introduce “Species by Proteome INvestigation” (SPIN), a proteomics workflow capable of querying over 150 mammalian species in 7.2 minutes of mass spectrometry (MS) analysis. Streamlined and automated sample preparation by protein aggregation capture, high-speed chromatography and data-independent acquisition, and a confident species inference algorithm facilitate processing hundreds of samples per day. We demonstrate the correct classification of known references, reproducible species identification in degraded Iron-Age material from Scandinavia, and test the limits of our methods with Middle and Upper Palaeolithic bones from Southern European sites with late Neanderthal occupation. While this initial study is focused on modern and archaeological mammalian bone, SPIN will be open and expandable with other biological tissues and taxa.

## Introduction

The potential of molecular species determination is generally unexploited when it comes to studies with large sample size, due to the prohibitive investments currently required for time-consuming analysis and data interpretation by experts. However, many discoveries can only be made from big data, such as following the change of an ecosystem^1^ and finding human bones in a “haystack” of faunal remains,^2^ or by continuous testing like food safety monitoring.^3^ Whenever the loss of anatomical features or heavy processing of biological tissues make morphological classification impossible, a decision needs to be made between two types of biochemical species identification: Targeted approaches like amplification-based polymerase chain reaction (PCR)^4^ and immunoassays^3^ can cope with enormous sample sets but can only detect one or a few species at a time. Global approaches, such as next-generation DNA sequencing (NGS)^5^ or proteomics,^6^ are theoretically able to identify any species, but the low throughput and high costs prohibit large studies, even with the latest nanopore sequencing technology.^7^ One method aimed at achieving both high throughput and global species identification is peptide-mass fingerprinting (PMF).^8, 9^ Unfortunately, it requires manual selection of species markers, mass spectra are mostly interpreted manually,^10, 11^ and the technology has become largely obsolete in proteomics^12^ because of lacking statistical control over peptide identifications^13^ and limited dynamic range without liquid chromatography (LC).

With the new “Species by Proteome INvestigation” (SPIN) workflow, we made use of the latest advances in LC-MS/MS to achieve fast and confident global species identification. Shotgun proteomics has the advantage of coping well with highly-degraded^14^ or processed^15^ specimens and for SPIN, we pushed the throughput beyond the fastest published methods^16^ and developed a universal species inference algorithm for reproducible data interpretation. We developed and benchmarked a single-step method to extract proteins from mineralized tissues, which is followed by automated contaminant removal and digestion by protein aggregation capture (PAC).^17, 18^ Shortening the LC-MS/MS analysis to less than 10 min became possible with new LC technology^19^ and fast-scanning data-dependent or multiplexed data-independent tandem MS acquisition methods.^20^ The new species inference algorithm based on a protein database with gene-wise alignment determines the most likely species and provides two quality control markers for estimating the confidence of each taxonomic assignment.

We decided to optimize and test the SPIN workflow for small 5 mg mammalian bone samples, as these are relevant for many potential use cases, such as forensic screening of animal or human remains,^21^ archaeological investigations of bones and tools,^22, 23^ authentication of processed food like gelatins,^15^ and analyzing the change of species diversity in ecosystems.^24, 25^ We optimized our methods with reference bones from 9 domesticated animals, a human, and 3 other great apes, validated the reproducibility and benchmarked comparability to morphology-based identification with more than 60 partially degraded bones from the Danish Early Iron-Age, and stress-tested the approach with a set of over 200 Middle and Upper Palaeolithic bone fragments from three archaeological sites in Portugal dating to 30 - 60,000 BP. We demonstrate that mammalian species families and taxa can be correctly assigned in most cases, while misassignments due to low signal intensity or gaps in the species sequence database can be avoided through thresholds based on the two quality control confidence scores.

## Results

### Streamlined and automated sample preparation

To increase throughput, we developed a new sample preparation protocol that consists of few manual steps allowing for easy scale-up (Fig. 1a). Changing the input material between finegrained bone powder and larger bone chips had only a minor impact on peptide identifications. We compared several different extraction buffer compositions and finally selected a mixture of hydrochloric acid and the non-ionic detergent NP-40 for combining demineralization and protein extraction, which are usually performed separately.^26^ This was the only tested combination that effectively prevented precipitation during the protein extraction and clean-up step. Contaminants, detergents, and minerals were removed using a modified protein aggregation capture (PAC) protocol, which allowed us to automate this step on a magnetic bead-handling robot using only 5 mg of bone material. We benchmarked our new sample preparation workflow against the commonly used “in-solution” digestion,^27^ “filter-aided sample preparation” (FASP),^28^ “gel-aided sample preparation” (GASP),^29^ and the more recent “S-trap”^30^ using the same Pleistocene Mammoth bone sample^31^ for all methods (Fig. 1b). The number of identified precursors by LC-MS/MS was the lowest for FASP and GASP, which was probably due to losses in the filter or gel. “In-solution” and “S-trap” performed about two-fold better, although “in-solution” had a relatively poor digestion efficiency leading to more missed tryptic cleavages. The new SPIN protocol produced significantly more precursor identifications and the automated sample preparation on a Kingfisher robot resulted in marginally fewer identifications but better reproducibility between the triplicates. From a practical standpoint, SPIN requires much less hands-on time than FASP, GASP, and “S-trap”, which cannot easily be scaled to 96-well format (data not shown). Although “in-solution” is similarly fast, the lack of protein clean-up makes it more susceptible to contamination problems, which complicates scale-up. With SPIN, a single laboratory worker can process four plates of bone chips (> 380 bone samples) per day using one magnetic bead-handling robot and four thermo-shakers, in parallel.

**Fig. 1.**
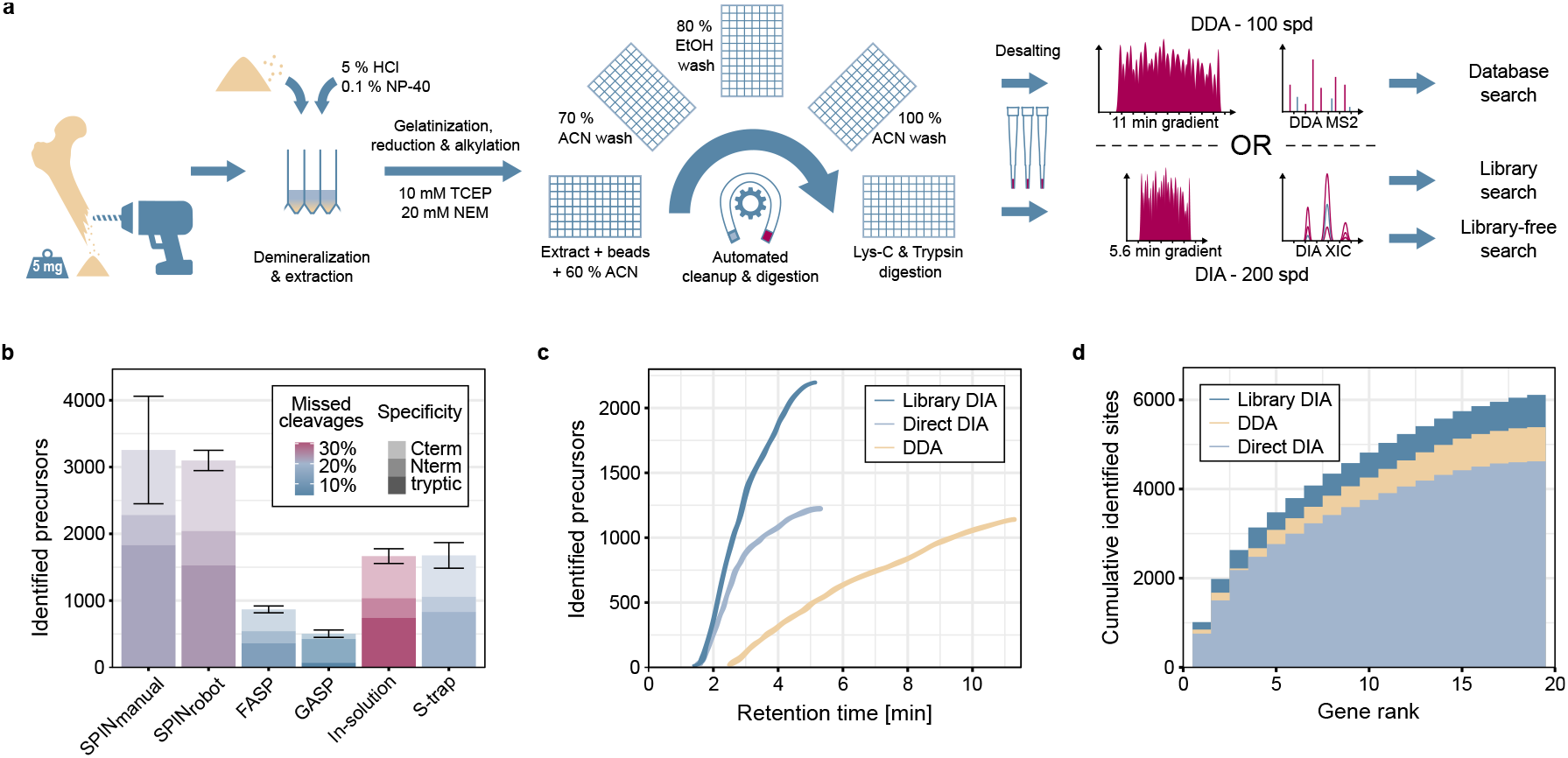
High-throughput bone proteome analysis workflow and benchmark. **a**, Sample preparation and data acquisition. Proteins are obtained from bone chips or powder by simultaneous demineralization and extraction. Cleanup by Protein Aggregation Capture and digestion can be automated by a magnetic bead-handling robot. Peptides are rapidly separated and analyzed by tandem mass spectrometry with a throughput of 100 samples per day (spd) in data-dependent or 200 spd in data-independent acquisition mode. **b**, Performance of the new “Species by Proteome INvestigation” (SPIN) protocol executed manually or by robot, compared to other common sample preparation techniques. Bone powder from a Pleistocene mammoth bone was used to perform each experiment in triplicates and peptides were analyzed with a 60 spd gradient and data-dependent acquisition. Mean unique precursor identifications are shown separately for enzyme specificity (lightness) and the average percentage of precursors with missed tryptic cleavage sites indicated by hue. Error bars indicate the standard deviation of total precursor identifications. **c**, Comparison of precursor identifications accumulated over retention time between the fast 100 spd DDA method analyzed by conventional database searching and the rapid 200 spd DIA method analyzed with a library-based vs. library-free approach. Peptides were generated by SPIN using a bovine bone and analyzed by LC-MS/MS with the two acquisition methods. **d**, Gene-wise cumulative absolute amino acid coverage based on the precursors identified in c) shown over the top 20 genes ranked by the number of precursors.

### Rapid peptide identification

SPIN uses very short online LC gradients to speed up the nanoflow chromatography coupled to Orbitrap tandem mass spectrometry. Compared to the throughput of the fastest acquisition methods in diagnostic plasma proteomics measuring about 50 samples per day^16^ or common methods in palaeoproteomics with less than 10 samples per day,^14^ we achieved sufficient proteome coverage for species discrimination of up to 200 samples per day with data-independent acquisition (DIA) and 100 samples per day with data-dependent acquisition (DDA).^19^ In a modern bovine bone sample, fast-scanning DDA identified about 1200 precursors in 11 min, while DIA reached the same number of identifications in 5.6 min when analyzed without a spectral library by directDIA (Fig. 1c). Almost twice as many precursors were identified by DIA analysis using a dedicated spectral library that needed to be generated once for each species of interest by DDA analysis of peptides offline fractionated by high pH reversed-phase chromatography.^32^ However, DIA yielded more redundant and overlapping precursors and therefore did not result in twice as much absolute sequence coverage (Fig. 1d). Across the 40 modern reference samples, the DDA method had a median coverage of 3678 amino acids and thereby outperformed the library-free directDIA approach with 3226 amino acids. The highest median coverage of 4480 amino acids was achieved with library-based DIA. As expected, sequence coverage was highest for the two most abundant genes, COL1A1 and COL1A2. Since there was almost no additional coverage to be gained beyond with more than 20 genes, we decided to exclude all other genes moving forward and thereby reduce noise and simplify database assembly.

### Species inference strategy

Completeness and quality of protein sequence databases is vastly different between taxa (Fig. 2b). Missing genes, gaps, and stretches of incorrect amino acid sequences can introduce a bias towards well-annotated species, if the phylogenetic assignment is done based on simple metrics like the number of identified peptides or protein groups. This is why we decided to build the species inference algorithm around site-based species comparison (Fig. 2a). It is based on a multiple sequence alignment (MSA) of each protein sequence of the 20 most common genes in bone for all available mammalian species. Further manual refinement is needed to remove faulty sequence inserts or errors like frame-shifts. To confirm that the sequence information of the 20 genes is sufficient for resolving the taxonomy of all 156 species in the database, we constructed a phylogenetic tree (Fig. 2b). The aligned protein sequence database is also the basis for creating a“site difference matrix” by performing a large species-to-species comparison, which contains only sites that are annotated but different within each pair-wise comparison. The aligned database is also used for mapping the LC-MS/MS-based peptide identifications to the correct genes and locations (“Mapping to alignment”, Fig. 2a). All precursors are combined into a normalized joined score (J-Score) for every detected sequence variant. Although the J-Score does not necessarily reflect the actual amino acid probability, it integrates multiple options and assigns them a weight. The summed J-Scores are used for determining the winner of every species-to-species comparison in the site difference matrix (Fig. 2c). There can be a single or multiple indistinguishable species winning most comparisons, which will be reported as the best matching species. We added an optional “fine grouping” step using a manually curated list of marker peptides to keep the phylogenetic placement between closely-related species consistent, even at low sequence coverage. To this point, a species will be assigned to every sample including the blanks. We established two control mechanisms for controlling and minimizing the false discovery rate (FDR) of our species calling algorithm, one to identify samples with too low signal, like blanks, and a second one to control for species that are not yet in the database. Samples with low peptide intensity are removed by measuring the abundance of autolysis-derived protease peptides, which we use like a spike-in standard. The threshold is calibrated based on relative protease intensity in laboratory blanks. The second control mechanism is aimed at the identification of species with insufficient sequence coverage, as this would lead to unreliable classification. Therefore, we extend the database with an equally sized set of decoy species (randomly-generated chimera species). The final results are then ranked by sequence coverage and a cutoff is applied to keep the number of decoy identifications below 1 %. The comparison of site coverage and relative protease abundance demonstrates that most of the blanks along with the empty samples were successfully removed using the two thresholds and that the coverage decreases with sample age (Fig. 2d).

**Fig. 2.**
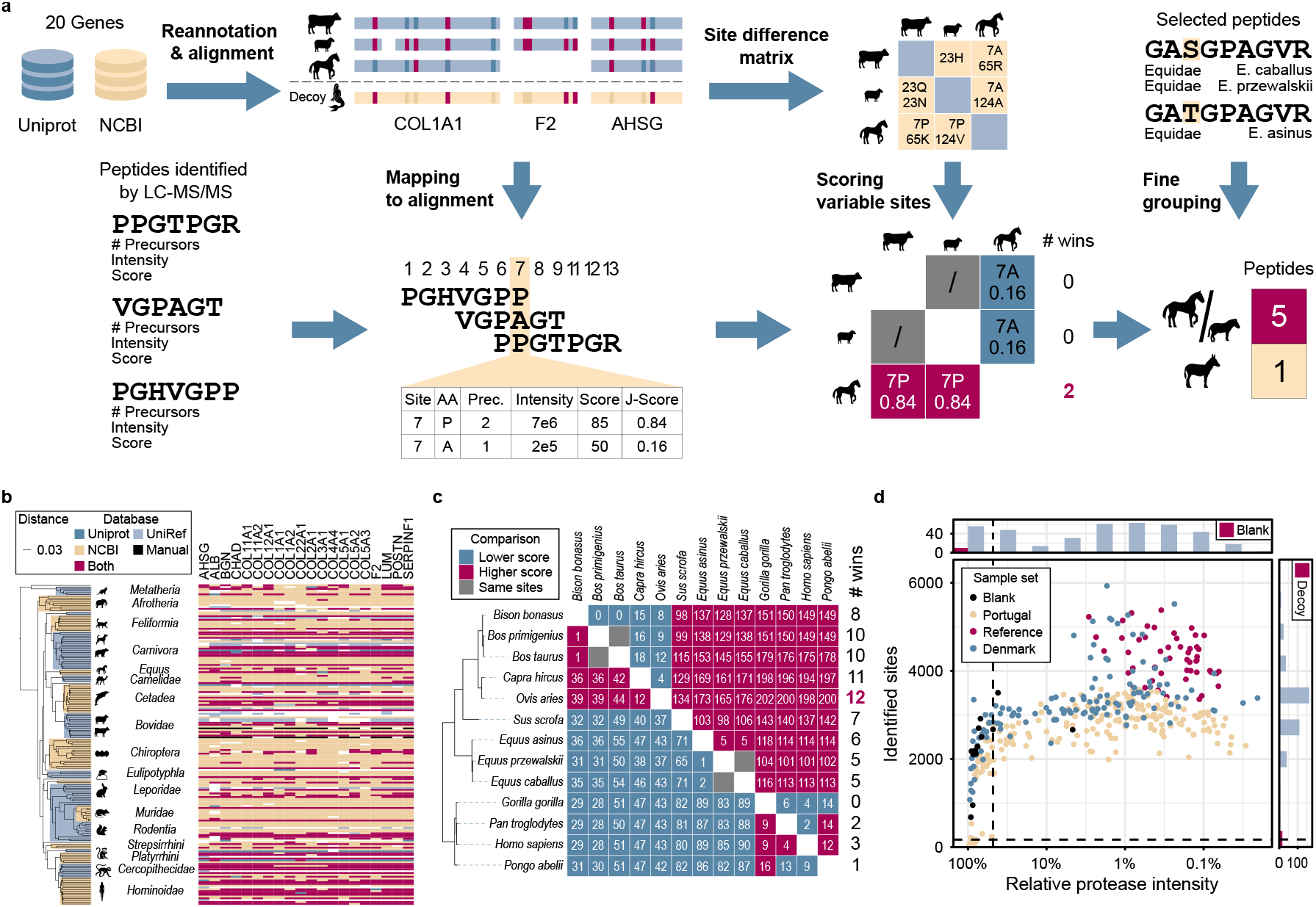
Data analysis pipeline for species identification. **a**, The aligned protein database, species difference matrix, and manually curated species marker peptides (top row) are used at multiple stages of the data processing pipeline (bottom row): Peptide identifications are converted to site level, scored by joining intensity, score, and number of precursors (J-Score) and used to identify the winner of all possible species-to-species comparisons. Fine resolution of closely-related species can be further improved by using manually selected species marker peptides. **b**, The mammalian species database comprising 20 genes across 177 species (156 species with > 14 genes) was generated by merging Uniprot and NCBI with manually curated and re-annotated protein sequences. The phylogenetic tree was generated from the protein database using Fast Tree and FigTree. **c**, Example species competition matrix for the reference sample “Ovis 07” only showing the 13 reference species. White numbers indicate the summed joint scores (J-Score) of the species-discriminating sites. Grey cells indicate species pairs, where no species-discriminating sites have been identified in the sample. Pink indicates that the left species wins and blue indicates that the top species wins the comparison. The phylogenetic tree is a subset of the tree in panel b. The complete species competition matrices comprise all 156 target and 156 decoy species, i.e. 24,336 comparisons. **d**, Absolute sequence coverage and relative protease intensity in reversed log10-scale for all samples from the three datasets in this study. The vertical site coverage cutoff is used to control the false-discovery rate at 1 %. The horizontal protease intensity cutoff excludes samples with low signal (lower than 75 % of the blank runs). Independent analysis of both parameters is displayed as histograms.

### Validating SPIN with bones from known species

We optimized and assessed the performance of the different data acquisition and interpretation strategies using a set of 49 known reference bones from 13 species (Fig. 3a). All samples were placed in the correct genus using library-based DIA, whereas library-free directDIA could not rule out human for chimpanzees, and DDA was not able to exclude goat for one of the eight sheep samples. Interestingly, all three methods performed equally well, when it came to the placement of taxa within the families. Within bovines, the domestic cattle (Bos taurus) could be distinguished from European bison (Bison bonasus) but not from the aurochs (Bos primigenius). The European bison itself could not be discriminated from American bison (Bison bison) and yak (Bos mutus) and in one case, from zebu (Bos taurus indicus) (Fig. 3b). The closely-related goat (Capra hircus) and sheep (Ovis aries), were correctly identified without the need for fine grouping in all DIA analyses and 10 out of 11 samples in DDA analysis.

**Fig. 3.**
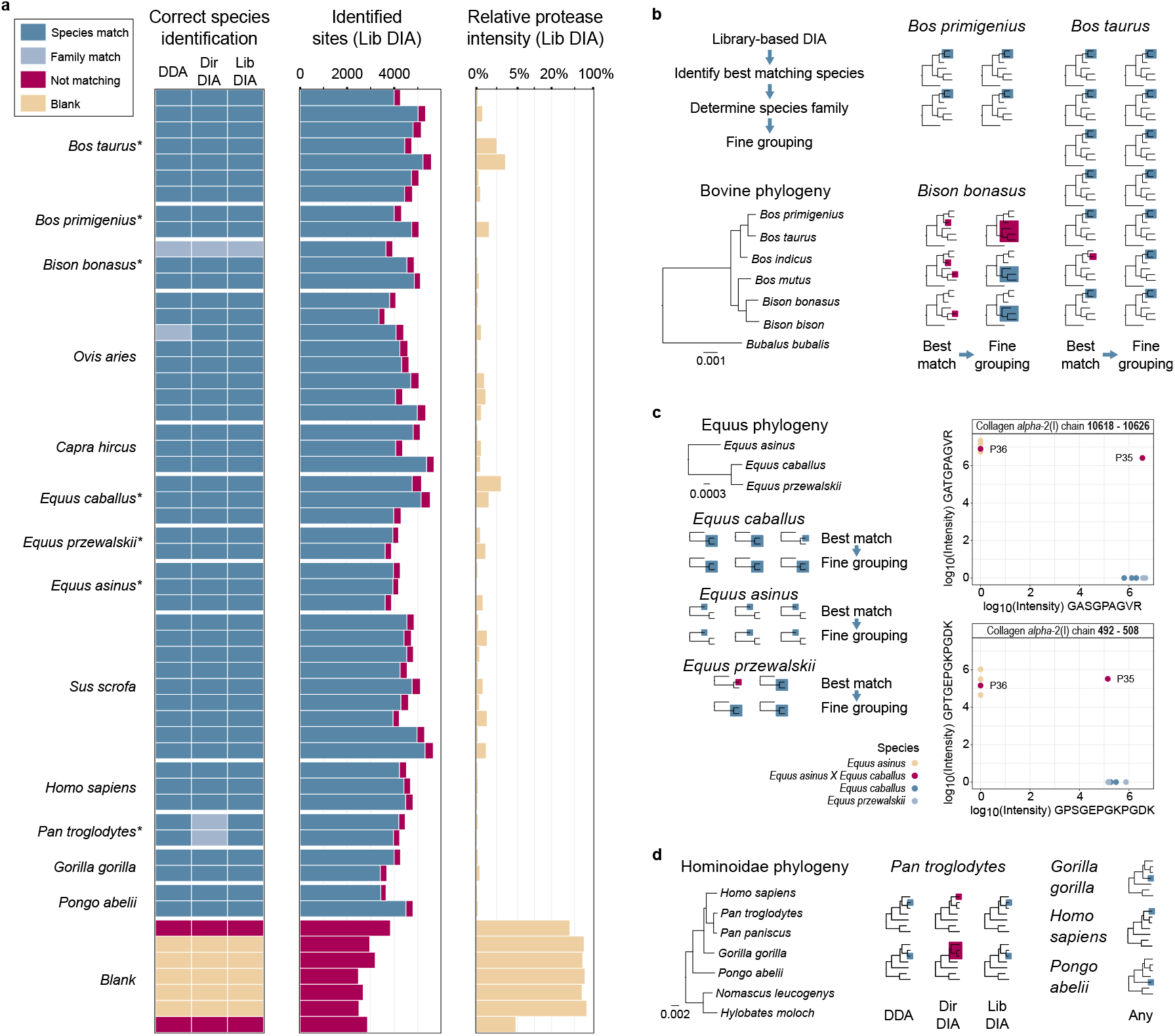
Reference species analysis. **a**, Species identification results based on DDA, directDIA, and library-based DIA analysis. Dark blue boxes indicate correct identification of a single or multiple indistinguishable (marked by asterisk) species. Light blue indicates species that could not be separated from their closest relatives. Blanks that were below the relative protease intensity threshold are shown in pink. The “identified sites” bar chart shows the absolute amino acid coverage in blue for sites matching the true species and pink for non-matching sites. The relative protease intensity is calculated by dividing the intensity of protease peptides by total intensity and plotted in log-scale. **b**, Bovine species identifications obtained by library-based DIA analysis. Phylogeny is based on the protein database. Correctly identified single or indistinguishable species are highlighted in blue. Inconsistent identifications are marked in pink. Best Matching species are on the left and the refined “fine-grouping” on the right side. **c**, Same display as in b for equine species analyzed by library-based DIA data. The additional plots on the right side show the log10 intensity of two species-discriminating peptides for the horse isoform on the x-axis and the donkey isoform on the y-axis. Missing quantifications are shown as zero log10 intensity. Hybrids are marked in pink. **d**, Species identifications after fine-grouping for great apes comparing the three different peptide identification strategies. Correctly identified single or indistinguishable species are highlighted in blue. Broader matches within the family are marked in pink. Differences between the three identification strategies were only observed for the genus *Pan*.

Within equines, domestic horse (Equus ferus caballus) was successfully discriminated from donkey (Equus africanus asinus), which cannot be done by PMF, but not from the Mongolian wild horse (Equus ferus przewalskii) (Fig. 3c). Fine grouping was not actually needed to exclude donkey, but it made the caballus/przewalskii classification more uniform. Besides common domesticated animals and their wild relatives, we explored the potential to detect and correctly identify great apes. While all three peptide identification methods could successfully classify human (Homo sapiens), orangutan (Pongo abelii), and gorilla (Gorilla gorilla), only DDA and library-based DIA analyses could correctly separate chimpanzee (Pan troglodytes) from Homo sapiens (Fig. 3d). It is noteworthy that both DDA and library-based DIA analysis of two chimpanzee bones assigned one of them to chimpanzee and the other to bonobo (Pan paniscus). Unfortunately, the low quality of the available bonobo protein database prevents closer investigation. These results confirm that the SPIN workflow can be used to classify great apes at genus level. As a proof-of-concept, we investigated the potential to detect species hybrids with SPIN by analyzing two samples from mules. Focussing on two peptides that are distinct between horse and donkey, one of the two mule samples showed high intensity for both sequence variants, as expected, but the second mule showed peptide intensities typical for a donkey (Fig. 3c). We concluded that SPIN is technically capable of hybrid detection, but an assessment of its reliability would require a larger study size.

### Performance and reproducibility benchmark

To benchmark the SPIN analysis strategy against standard-practise bioarchaeological species determination based on bone morphology, we analyzed a set of 64 bone fragments related to human activities at the “Salpetermosen Syd 10” site (MNS50010, ZMK5/2013) in Denmark, which dates to the early Pre-Roman Iron-Age site (380 BC - 540 AD, Fig. 4a).^33^ Some of the bones showed strong signs of decay due to the age and the wet anoxic conditions in the Salpetermosen bog. Each specimen was morphologically analyzed by an experienced zooarchaeologist and the SPIN analysis was conducted in duplicates with high and low MS loading amounts. The study was blinded by keeping the morphological species identification undisclosed, until the SPIN analysis was finalized.

**Fig. 4.**
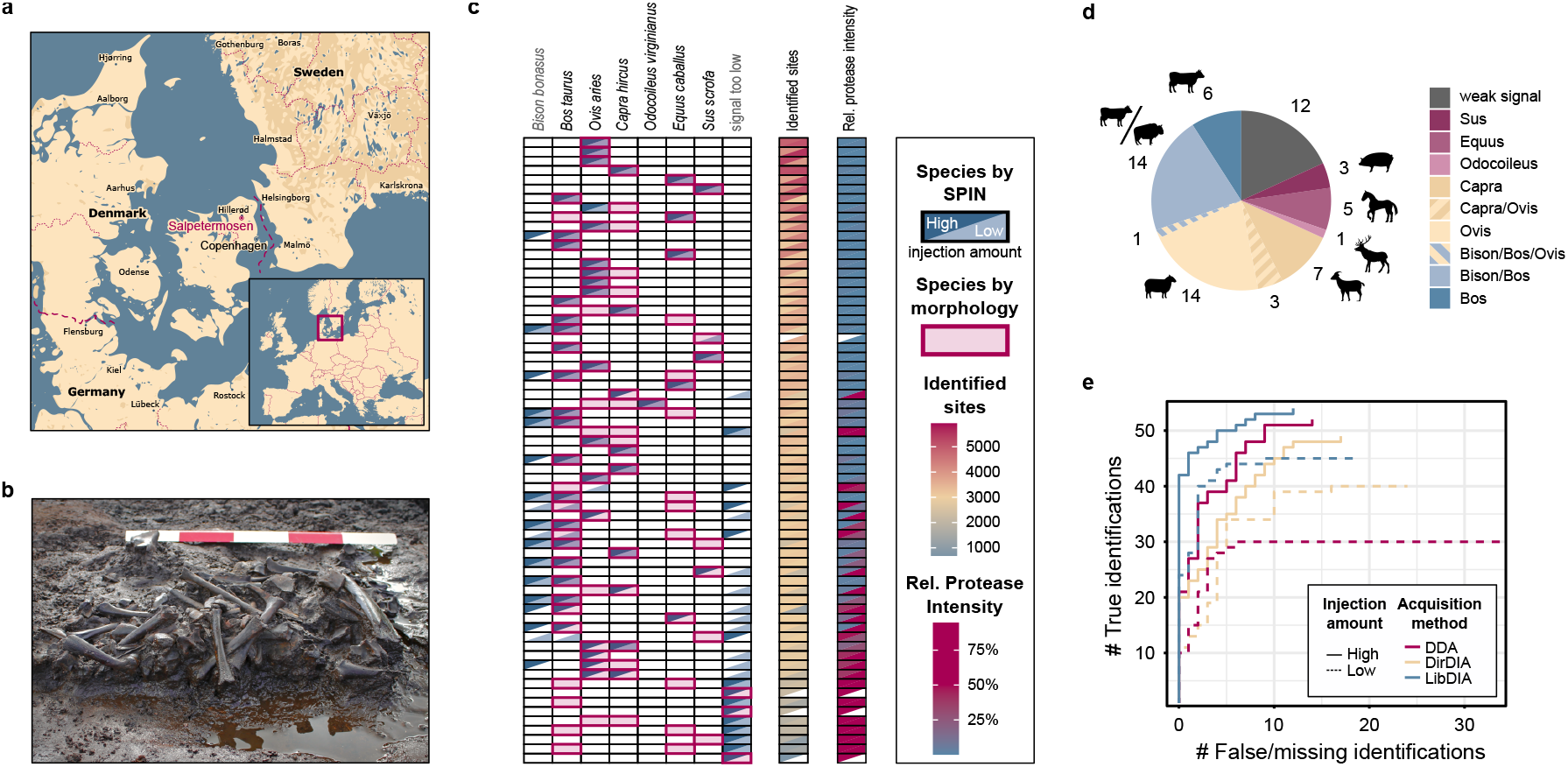
Species identification of bones from the Scandinavian Iron-Age. **a**, Location of the archaeological site “Salpetermosen Syd 10” on Zealand in Denmark in the Hillerød municipality 30 km north of Copenhagen. Map drawn in Mapbox Studio using a custom style. **b**, Cross section of an in situ wetland bone deposit. Scale bar is 50 cm. Four bones were radiocarbon dated between 1,720 - 1,570 BP. Picture provided by the Museum of North Zealand. **c**, Individual species identification results of all 63 samples and 3 blanks. The upper left and lower right wedge of each cell represent replicate results measured with high (upper left) and low (lower right) MS injection amounts. The first matrix shows species identified by SPIN in blue and pink boxes indicating morphological species identifications from a zooarchaeology expert. The second and third matrix are heatmaps showing the absolute number of covered amino acids and relative protease intensity, respectively. **d**, Overall species identifications obtained with library-based DIA and fine-grouping. Striped colors indicate samples with insufficient sequence coverage to distinguish closely-related taxa. The group “weak signal” contains samples below the relative protease intensity threshold including the 3 blanks. **e**, Pseudo receiver operating characteristic (ROC) curves for comparing the sensitivity and success-rate of three different data-acquisition and analysis strategies. Results of each dataset were sorted by decreasing number of identified sites. The y-axis shows the cumulative number of correct species identifications in agreement with the morphology. The x-axis shows the cumulative number of false or missing identifications below the relative protease intensity threshold.

SPIN analysis with library DIA and fine grouping resulted in 49 exact and 3 approximate species identifications, whereas 11 samples were excluded due to the low peptide intensity (Fig. 4b). The comparison of replica analyses showed perfect reproducibility between duplicates with site coverage ¿ 3000 amino acids (Fig. 4c). With lower coverage, more samples - mostly the low injection replicates - stayed below the relative protease intensity cutoff. There was only one case of contradicting species identifications be tween the two replicates. In that case, cattle was identified in the “low” and horse in the “high” replicate, while morphology confirmed it as horse.

For 96 out of 99 identified samples from both replicates, the SPIN analysis by library-based DIA was in agreement with the morphological analysis (Fig. 4c). Sheep and goat, which often cannot be discriminated morphologically, were unambiguously identified by SPIN in 21 cases and could not be distinguished in two. For cattle, only Bos is plausible at this time and location^34^ and the SPIN and morphological identifications showed good agreement. Since the SPIN algorithm does not know the archaeological context, Bison was still considered and we observed that discriminating between Bos and Bison became significantly more challenging with lower sequence coverage. Cattle and horses could not be distinguished morphologically in 9 cases, 6 of which could be resolved by SPIN. All three laboratory blanks (marker at “signal too low”) were correctly excluded by the relative protease intensity threshold.

We compared the performance of the three dif ferent types of peptide identifications by library DIA, directDIA, and DDA, which performed very similarly for the reference samples. The differences became much more apparent in the Salpetermosen sample set. The pseudo ROC-curve analysis shows that the DIA-based methods outcompeted DDA, especially in the low amount replicate indicating higher sensitivity in DIA-based measurements (Fig. 4d). Between the two DIA methods, library-based DIA consistently produced more true species identifications than directDIA.

### Species identification of Middle and Upper Palaeolithic archaeological bones

To challenge the SPIN workflow with highly-degraded samples and demonstrate its scalability, we analyzed a set of 213 archaeological bone fragments from three Portuguese archaeological sites with early human occupation (Fig. 5a). To this end, we translated the output of the species inference algorithm to reflect the most likely ancestors that were present at the location and time. Analogous to all other samples in this study, we analyzed the Portuguese bones with DDA and DIA, which took 52 h and 26 h of MS acquisition time, respectively. We used the library-based DIA results as the basis for species identification because of its higher resolution, as demonstrated with the Salpetermosen dataset. However, to allow the identification of species for which no spectral library is currently available, such as rodents, we replaced the result with the taxonomy identified by directDIA, whenever directDIA detected such a species. As both results are based on the same raw data, the relative protease threshold remains unaffected, but the sequence coverage is lower with direct-DIA. In addition, compared to the reference and Salpetermosen samples, these Southern European Middle and Upper Palaeolithic samples suffer from reduced protein sequence coverage across the proteome assembly (Fig. 5c).

**Fig. 5.**
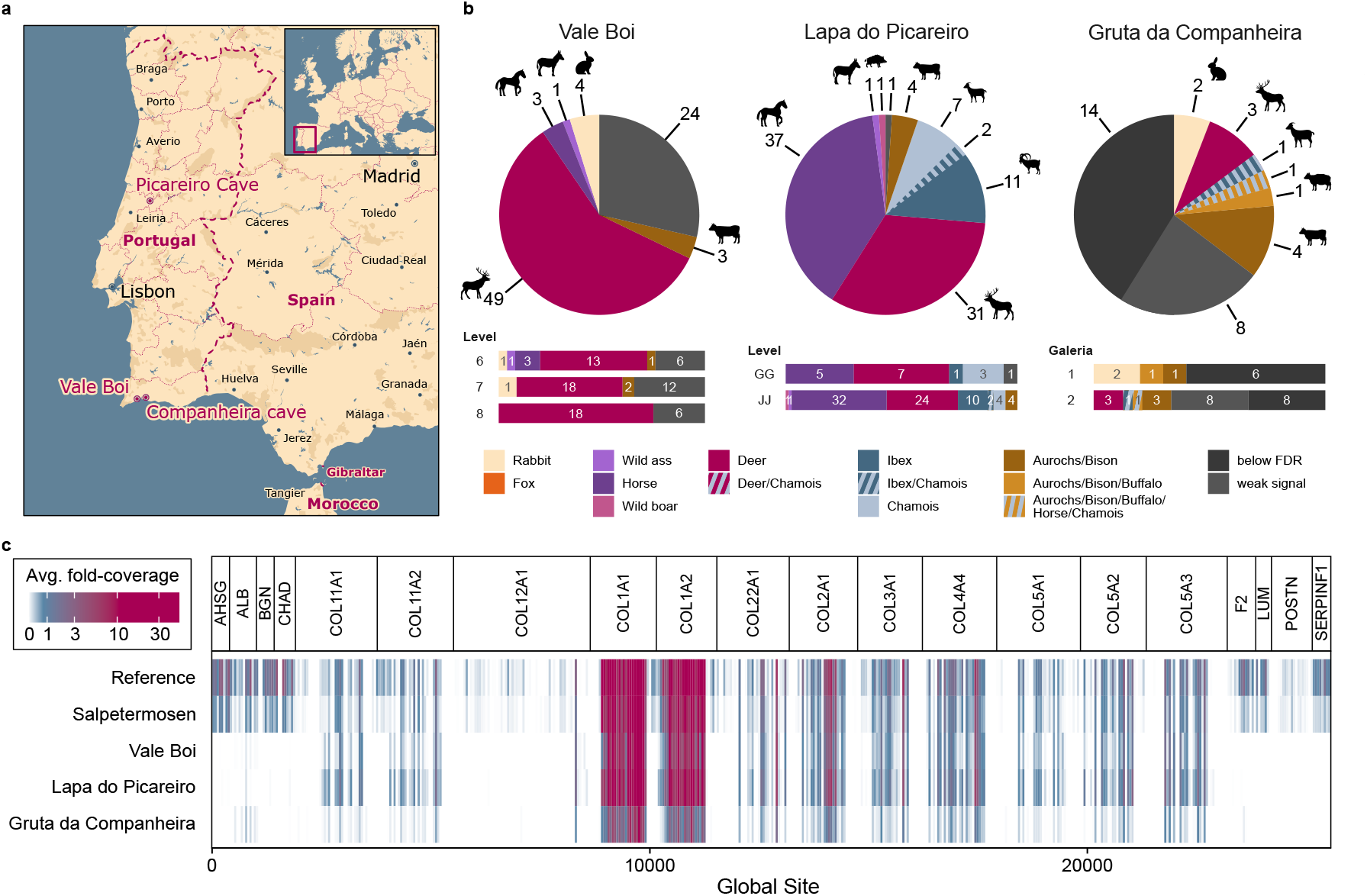
Large-scale species identification at 3 sites with early human occupation on the Iberian peninsula. **a**, Locations of the three sites on a current map of Portugal. Map drawn in Mapbox Studio using a custom style. **b**, Species identified in 84 samples from levels 6 - 7 (29 - 31,500 BP) of Vale Boi, in 95 samples from layers GG to JJ (38 - 45,000 BP) of Lapa do Picareiro, and 34 samples from chambers 1 and 2 (estimated 50 - 60,000 BP) of Gruta da Companheira. **c**, Average fold-coverage of the 20 genes used for SPIN comparing the 3 Portuguese sites with the modern reference and iron age material. Coverage is calculated by summing the number of precursors at each site in the global aligned database. The values represent the average fold-coverage in 10 amino acid bins for each data set.

For Lapa do Picareiro, 94 out of 95 samples could be assigned a confident species identification. For these specimens, that date approximately between 38,000 - 41,000 BP (layers GG-II) and 45,000 BP (layer JJ),^35, 36^ species composition is relatively similar for both layers (Figure 5b). Of particular interest is the identification of one specimen of the now-extinct European wild ass (E. hemionus hydruntinus), alongside 37 caballine horses. Most of the ibex and chamois bones, which are not easy to distinguish morphologically, could be uniquely assigned to one of the two (18 out of 20). Finally, bovines and wild boar were exclusively identified in the older “JJ” layer. For Vale Boi, dated between 31,500 and 29,000 BP,^37^,^38^ 60 out of 84 samples could be assigned a confident species identification. The remaining 24 samples failed to meet the abundance-based quality threshold and were therefore not assigned a species identity. The vast majority of the identified bones were classified as deer, which is in agreement with the previously reported numbers for large mammals.^37^ Equids, including one E. hydruntinus, were only identified in layer 6. With direct-DIA, four smaller bone fragments from layers 6 and 7 could be classified as rabbits, which were highly abundant at Vale Boi.^39^

Finally, for Gruta da Companheira 20 out of 34 samples could be assigned a confident species identification. Expected to date around 50,000 to 60,000 BP, 14 samples failed to meet the FDR threshold while eight samples were excluded due to failing to meet the abundance-based quality threshold. Amongst the confidently identified species at Gruta da Companheira were bovines, deer, and rabbits, whereas the only ovicaprine sample could not be uniquely assigned to either ibex or chamois. Although below the relative protease cutoff, the two most interesting bones, which were both found in Galeria 2, matched best to great apes. The sample with highest sequence coverage (1817 aa) was classified as human or chimpanzee, whereas the sample with lower coverage (235 aa) matched equally well to all great apes. Here, the SPIN results can be used as a starting point for future indepth protein and ancient DNA analyses to find out whether these are actually human remains, and to eventually define their genetic profile.^14^

## Discussion

Here, we present a new workflow for genetic species identification that bridges the gap between high-throughput targeted methods, such as PMF, PCR, or ELISA, and more powerful low-throughput approaches like NGS or conventional proteomics analysis by LC-MS/MS. We demonstrate that the automated sample preparation workflow based on PAC produces high quality peptide samples for bone proteome analysis by LC-MS/MS and allows for easy scale-up. Data acquisition, which used to be a bottle neck in terms of speed and costs, was drastically shortened to reach a throughput of up to 200 samples per day per MS instrument. This is possible with a new gradient storage-based LC system and advanced software for DIA data interpretation, two emerging technologies that are undescribed in bone proteome analysis. The data interpretation strategy used for SPIN facilitates an unbiased comparison of currently up to 156 species and provides high confidence due to FDR control on both peptide and species identification level. We concluded that the highest confidence and sensitivity could be achieved with the library-based DIA approach, based on the analysis of 64 partially degraded bones from Denmark. These results were validated internally with replication and externally with morphological species identification.

Furthermore, we demonstrated the high-throughput of this method by analyzing a set of 213 Palaeolithic bones from Portugal in little more than a single day of MS time. SPIN performed well during the analysis of degraded samples and still had a reasonable success-rate for 50,000 - 60,000 year-old bone material from mediteranean climate with hot, dry summers and cool, wet winters. In contrast, samples of similar thermal age require considerable effort for ancient DNA analysis. We suppose, with an optimized sample preparation protocol for very stable specimens like teeth, SPIN will become applicable for palaeoproteomics analysis of million-year-old material.^14^

So far, the SPIN data analysis is limited to 156 species and 13 spectral libraries for improved library-DIA analysis. We envision that these numbers will grow with more research groups sharing their protein sequence databases and spectral libraries. Furthermore, the SPIN workflow itself will likely be improved and expanded over time. We think of it as a modular protocol that can serve as the foundation for “SPIN-off” methods with custom building blocks, like: (i) sample preparation modified to support protein extraction from heavily-processed food products,(ii) data acquisition adapted for different instruments, or (iii) data interpretation including sex identification. We already touched upon more advanced use cases for SPIN like the identification of hybrid species, such as mules. Furthermore, it can be adapted, to resolve mixtures of proteins from multiple taxa or to quantify protein damage, in the future. Finally, we anticipate that the SPIN workflow will make LC-MS/ MS more accessible for everyone, due to the reduction of the analytical costs per sample and high degree of automatization.

## Methods

### Sample description

Each bone sample in this study was taken with the permission from the respective museum, curator, or institution and the impact was minimized by only removing necessary amounts. A fragment of a Pleistocene mammoth bone from permafrost and dated to approx. 43,000 BP^31, 40^ was used for optimizing methods. Reference samples for Bos taurus, Ovis aries, Sus scrofa (mandibles), and Equus caballus (phalanx) were from the mixed viking-medieval deposits of the archaeological site Hotel Skandinavien (Århus Søndervold) (ZMK139/1964) in Århus, Denmark. The Department of Forensics at the University of Copenhagen provided the human reference sample, dentine from a previously described^14^ 200 - 400 year-old premolar from “Almindelig Hospitals cemetery on Østerbrogade” in Copenhagen, Denmark. Reference samples for Bos primigenius, Bison bonasus, Capra hircus, Equus asinus, Equus primigenius, mule (Equus caballus X Equus asinus), Pongo pygmaeus, Gorilla gorilla, and Pan troglodytes were provided by the Natural History Museum of Denmark. The set of 213 upper-palaeolithic bone fragments from three Portuguese sites consisted of 84 samples from level 6 (29,300 BP^37^), level 7 (30,400 BP^38^), and level 8 (31,500 BP, unpublished) from Vale Boi, 95 samples from layers GG-II (38.000 - 41,000 BP) and JJ (45,000 BP^35, 36^) from Lapa do Picareiro, and 34 samples from Galeria 1 and 2 from Gruta da Companheira.^41^

### Sampling

The reference samples for Bos taurus, Ovis aries, Sus scrofa, Equus caballus, and Homo sapiens and the 213 Portuguese samples were sampled in a clean laboratory designed for ancient DNA and protein work at the GLOBE Institute, at the University of Copenhagen. The remaining samples were processed in a laboratory with measures against human protein contamination. Working areas and tools were decontaminated with 5 % bleach and 70 %ethanol, between samples. Reference bones intended for the generation of spectral libraries were surface-decontaminated by mechanical ablation and small samples of ¡ 100 mg were removed using a rotary cutting tool, followed by crushing of the pieces by mortar and pestle. For high-throughput analysis, approx. 5 mg samples were collected by scraping the fracture site of bone fragments with a small chisel or pliers and transferred into a 96-well plate. At least 3 laboratory blanks, i.e. empty wells, are included for each project with at least 1 blank on every plate.

### Combined demineralization and extraction for SPIN

Five milligrams of bone powder or chips are suspended in 100 *μ*l 5 % HCl and 0.1 % NP-40 (ThermoFisher Scientific, 28324) in ultrapure water. Demineralization takes place at room temperature (rt) with continuous shaking at 1000 rpm, for 16 - 24 h. Reduction, alkylation, and collagen gelatinization are facilitated by adding 10 *μ*l 0.1 M tris(2-carboxyethyl)phosphine (TCEP, Sigma-Aldrich, C4706) and 0.2 M N-ethylmaleimide (NEM, Sigma-Aldrich, E3876) in 50 % ethanol and 50 % ultrapure water and shaking at 1000 rpm at 60 °C, for 1 h.

### Protein cleanup and digestion for SPIN

The purification and digestion take place on a KingFisher™ Flex (ThermoFisher Scientific) magnetic bead-handling robot. Debris is removed from the protein extract by centrifuging the plate at 800 x g, for 5 min. Magnetic SiMAG-Sulfon beads (Chemicell, 1202) are washed and prepared at a final concentration of 5 mg/ml in 60 % acetonitrile (ACN) 40 % water. In a deep-well KingFisher™ plate, 10 *μ*l bead solution and 40 *μ*l of the clear protein extract are briefly mixed. Protein aggregation capture (PAC) is initiated by the addition of 240 *μ*l 70 % ACN and 30 % water (60 % final ACN concentration) and finalized by incubating with shaking at 800 rpm for 5 min and without shaking for 1 min. The robot is loaded with this plate, “wash I” (500 *μ*l 70 % acetonitrile, 30 % water), “wash II” (500 *μ*l 80 % ethanol, 20 % water), “wash III” (500 *μ*l 100 % acetonitrile, and the “on-bead-digestion” plate (100 *μ*l 20 mM Tris pH 8.5, Sigma-Aldrich 10708976001, 1 *μ*g/mL LysC, Wako 129-02541, 2 *μ*g/mL Trypsin, Promega V5111). The programmed sequence is: (i) collect the beads with low speed for 3:30 min, (ii - iv) washes I-III with slow mixing for 2 min, (v) digestion at 37 °C with slow mixing for 1 h, and (vi) bead collection and removal. The digestion is finalized outside the robot with shaking at 800 rpm and 37 °C, overnight. The peptides are acidified with 10 *μ*l 10 % trifluoroacetic acid (TFA, Sigma-Aldrich, T6508). One Evotip (Evosep, EV-2001) per sample is washed in ACN, soaked with isopropyl alcohol, and equilibrated with 0.1 % TFA in water, according to the manufacturer’s protocol. The equilibrated tips are loaded with 10 *μ*l peptide solution and subsequently washed with 20 *μ*l 0.1 % TFA, before LC-MS/MS.

### Peptide fractionation for spectral libraries

Peptides for spectral libraries were obtained either from 3 x 5 mg bone powder processed by robot-based SPIN or from 20 mg bone powder processed manually with the same workflow. The peptides were desalted using C18 (3M Empore, 66883-U) StageTips.^42^ After quantification by Nanodrop (Thermo Fisher Scientific) at A280nm, 12 *μ*g peptides were adjusted to pH 7-8 by adding one volume of 50 mM ammonium bicarbonate (ABC, Sigma-Aldrich, A6141). Offline fractionation by high-pH reversed-phase chromatography was carried out on an Ultimate 3000 HPLC (Thermo Fisher Scientific) equipped with a 15 cm long, 1 mm i.d., 1.7 *μ*m particle size C18 column (Waters ACQUITY Peptide CSH) and 5 mM ABC as buffer A and 100 %ACN as buffer B. The gradient at a flow rate of 30 *μ*l/min started at 6 % B, was increased to 18 % B over 55 min, to 25 % B over 12 min, and to 70 % B over 3 min, followed by a column wash at 70 % B for 7 min and re-equilibration at 6 % B for 9 min. During the gradient, 12 fractions of equal size were collected. Blanks were run between the different species to reduce carryover. The fractions were acidified with 1 % (final concentration) TFA and vacuum concentrated. An equivalent of 250 ng peptides (25 % of each fraction) was loaded on Evotips as described above.

### Protein extraction and digestion methods for comparison

Each of the tested protocols “in-solution”,^43^ FASP,^44^ GASP,^29^ and S-trap^30^ was conducted in triplicates using 5 mg bone powder from the Pleistocene mammoth bone test sample. Demineralization was done by adding 100 *μ*l 5 % HCl and incubating at r.t. and shaking at 1000 rpm, for 24 h. Insolubles were separated by spinning the suspension at 5,000 x g for 5 min and the supernatant was discarded or kept for FASP. The protein pellet was washed with 100 *μ*L ultrapure water followed by repeating the centrifugation and discarding the supernatant. Proteins were extracted either in 100 *μ*l 3 M guanidinium hydrochloride (Gnd-HCl, Sigma-Aldrich, G3272) in 0.1 M Tris, pH 8.5 for “in-solution” and FASP or in 100 *μ*l 2 % SDS (Sigma-Aldrich, 428018) for GASP and S-trap. To all protein extractions, 10 mM TCEP and 20 mM chloroacetamide (CAA, Sigma-Aldrich, C0267) were added and the samples were incubated with shaking at 1000 rpm at 60 °C, for 1 h. The protein concentrations of the extracts were quantified by BCA assay (ThermoFisher Scientific, 23225). For “in-solution”, the guanidinium extract was incubated with 1:300 (protease_wt_:protein_wt_) LysC at 37 °C, for 1h. Subsequently, the sample was diluted with 3 volumes of 25 mM Tris, pH 8.5 and 1:100 (protease_wt_:protein_wt_) trypsin were added for an overnight digestion at 37 °C. The protein digest was acidified with 1 % (final concentration) TFA and desalted using StageTips.

For FASP, the guanidinium extract was spun at 20,000 x g, for 10 min and the supernatant was mixed with 9 volumes 8M urea in 0.1 M Tris, pH 8.5 and transferred and passed through a 2.5 kDA MWCO filter (Millipore, UFC500324) in 200 *μ*l steps by spinning at 5,000 x g for about 10 min. The filter was washed with 200 *μ*l 8M urea in 0.1 M Tris, pH 8.5 the demineralization supernatant was mixed with 6 volumes 8M urea in 0.1 M Tris, pH 8.5 and passed through the same filter in 200 *μ*L steps. After the last wash with 200 *μ*L 8M urea in 0.1 M Tris, pH 8.5, 100 ng LysC in 100 *μ*l 50 mM ABC were added to the filter. Pre-digestion took place at 37 °C with shaking at 1000 rpm, for 1h. Next, 200 ng trypsin were added and the digestion continued at 37 °C, overnight. The peptides were collected by spinning the filter and washing the membrane with 100 *μ*l Tris, pH 8.5, spinning, washing with 100 *μ*l 40 % ACN 60 % water, and spinning for final collection. The flow-through was concentrated to ¡ 50 *μ*l and acidified with 1 % (final concentration) TFA, before desalting on C18 StageTips.

For GASP, 100 *μ*l SDS extract was mixed with 100 *μ*l acrylamide solution (37.5:1 Acrylamide:Bisacrylamide), for 20 min. Polymerization was started by adding 8 *μ*l TEMED (Sigma-Aldrich, T9281) and 8 *μ*l 10 % ammonium persulfate (Sigma-Aldrich, A3678). The obtained gel was sliced into pieces and fixed in 50 % methanol, 40 % water, 10 % acetic acid, for 30 min. The supernatant was removed and the gel was sequentially washed with 1 ml ACN, 1 ml 6 M urea, 1 mL ACN, 1 mL ACN, 1 ml 6 M urea, 1 mL ACN, 1 mL 50 mM ABC, 1 mL ACN, and finally resuspended in 200 *μ*l 50 mM ABC. For predigestion, 100 ng LysC were added and the sample incubated at 37 °C with shaking at 1000 rpm, for 1h. Next, 200 ng trypsin were added and the digestion continued at 37 °C, overnight. The supernatant containing the peptides was collected and the gel pieces extracted using 200 *μ*l ACN, followed by 200 *μ*l 5 % formic acid, and 200 *μ*l ACN. The peptide solutions were combined, vacuum-concentrated, and desalted on C18 StageTips.

For S-trap, 1.2 % (final concentration) phosphoric acid was added to the SDS extract. An equivalent of 10 *μ*g extract were mixed with 1 volume 90 % methanol, 100mM triethylammonium bicarbonate, pH 7 (TEAB, Sigma-Aldrich, 18597) and loaded on an S-trap filter (Protifi) by spinning at 2,000 x g, for 1 min. The flow-through was re-loaded three times, before washing the filter with 200 *μ*L 90 % methanol, 100mM TEAB, pH 7. For predigestion, 100 ng LysC in 100 *μ*l 50 mM Tris, pH 8.5 were added and the samples incubated with shaking at 1000 rpm at 37 °C, for 1 h. Then, 200 ng Trypsin in 50 *μ*l 50 mM Tris, pH 8.5 were added and the digestion continued, overnight. The peptides were collected by centrifugation at 2,000 x g, for 1 min, and sequential washing with 100 *μ*l 50 mM Tris, pH 8.5, 100 *μ*l 0.1 % TFA, and 100 *μ*l 0.1 % TFA in 50 % ACN. After concentrating the peptides by vacuum evaporation, they were desalted on C18 StageTips.

### Data acquisition methods

For SPIN by DDA, chromatography was carried out using the 100 samples per day (SPD) method of an Evosep One (Evosep, Odense, Denmark) and an analytical column made in-house using a laser-pulled 8 cm long 150 *μ*m inner diameter capillary packed with 1.9 *μ*m C18 particles (Reprosil, Dr. Maisch). Peptides were ionized by nano-electrospray at 2 kV and analyzed on an Orbitrap Exploris 480TM (Thermo Fisher Scientific, Bremen, Germany) MS. Full scans ranging from 350 to 1400 m/z were measured at 60 k resolution, 25 ms max. IT, 300 % AGC target. The top 6 precursors were selected (30 s dynamic exclusion) for HCD fragmentation with an isolation window of 1.3 m/z and an NCE of 30. MS2 scans were acquired at 15 k resolution, 22 ms max. IT, and 200 % AGC target.

DIA SPIN analysis was based on the 200 SPD method of an Evosep One and the same column, ESI, and MS instrument as for DDA. Full scans ranging from 350 to 1400 m/z were measured at 120 k resolution, 45 ms max. IT, 300 % AGC target. Precursors were selected for data-independent fragmentation in 15 windows ranging from 349.5 to 770.5 m/z and 3 windows ranging from 769.5 - 977.5 m/z with 1 m/z overlap. HCD fragmentation was set to an NCE of 27 and MS2 scans were acquired at 30 k resolution, 45 ms max. IT, and 1000 % AGC target. Samples for method optimization and spectral libraries were measured in DDA mode. Method optimization experiments were analyzed on the 60 SPD Evosep One gradient and spectral libraries on the 200 SPD gradient using the same column, ESI, and MS instrument as described above. Full scans ranging from 350 to 1400 m/z were measured at 60 k resolution, 25 ms max. IT, 300 % AGC target. The top 12 precursors were selected (30 s dynamic exclusion) for HCD fragmentation with an isolation window of 1.3 m/z and an NCE of 30. MS2 scans were acquired at 15 k resolution, 22 ms max. IT, and 200 % AGC target.

### Protein database for SPIN

The database includes all protein sequences on UniProt knowledgebase^45^ and NCBI RefSeq^45, 46^ from mammalian species and matching to the top 20 genes (Fig. 2b) in .fasta format. The NCBI entries were re-annotated with gene and species information using the respective GenPept files and the fasta-headers were changed to a pseudo-Uniprot format: “>NCBI [protein ID] [protein ID] [gene alias] [protein description] OS=[species name] OX=[species ID] GN=[gene name]”. Relevant Uniprot entries with missing or false gene annotations were added by sequence similarity-based re-annotation. The UniRef90 (release 2020 06) repository was used to annotate each “90 % similarity” cluster with its most common gene name, followed by downloading all new proteins matching the top 20 genes and updating the fasta-headers to include the correct gene name. Protein sequences from species missing in the databases like Equus przewalskii, Bison bonasus, and the extinct Bos primigenius were manually extracted from the available genomes^47–49^ using reference sequences of the closest living relatives and the local BLAST^50^ and visualization in UGENE.^51^ After combining all protein sequences, filtering for mammalian species, and removing duplicates, the sequences were split into 20 separate files for each gene.

A multiple sequence alignment (MSA) was performed for each file using MUSCLE version 3.8.425^52^ and the alignment was visualized in AliView version 1.26.^53^ Upon manual inspection, faulty sequences, for instance large inserts that were not shared by any other species or frameshifts identified by very low similarity to the rest of the alignment, were removed or changed into gaps. The aligned and manually refined databases were combined into one .fasta file with gene-wise alignment and a second gapless file for the use in search engines. Phylogenetic trees based on this database were made by merging all genes of each species in alphabetical order, generating a consensus, generating the tree with FastTree version 2.1.11,^54^ and visualization with FigTree version 1.4.4.^55^

### Peptide identification

All raw files from DDA were analyzed in MaxQuant version 1.6.0.17.^56^ Variable modifications included oxidation (M), deamidation (NQ), Gln −> pyro-Glu, Glu −> pyro-Glu, and proline hydroxylation, whereas NEM-derivatization of Cys was configured as fixed modification. All files searched against a complete protein database, such as searches of spectral libraries of method optimization runs, were first run with tryptic specificity and up to 2 missed cleavages to reduce the database size and then searched again with semi-tryptic specificity allowing for a peptide length between 8 - 30 aa and max. mass of 4000 Da. SPIN files were searched against the above mentioned gapless database only with semi-tryptic specificity using the same settings. Up to 5 variable modifications were allowed in tryptic and up to 4 in semi-tryptic searches. In all searches, “Second peptides” search and “Match between runs” were disabled, the score thresholds for identification were set to a minimum Andromeda score of 40 and delta score of 6, and the internal MaxQuant contaminant list was replaced with a custom database. All other settings were left as default.

All raw files from DIA were analyzed in Spectronaut version 14.5.200813 (Biognosys)^57, 58^ using either library-based or library-free DirectDIA search. Spectral libraries were imported from the individual semi-tryptic MaxQuant search results with default settings except digestion-specificity and merged into a single library, before searching. The library-based search was carried out with default settings, except for MS1 as “Quantity MS-Level”. The DirectDIA search was configured with the abovementioned gapless and custom contaminant databases. Digestion specificity was set to semi-tryptic with a peptide length between 7 - 40 amino acids, up to 2 missed cleavages, and the variable modifications included oxidation (M), deamidation (NQ), Gln −> pyro-Glu, Glu −> pyro-Glu, and proline hydroxylation, whereas NEM-derivatization of Cys was configured as fixed modification. Again, the “Quantity MS-Level” was set to MS1 and all other settings were kept as default.

### Species inference

The species determination based on peptides identified with library-based DIA, DirectDIA, or DDA was done in R version 4.0.3 using RStudio version 1.3.10.93 and additional packages. The required peptide identification data is either generated with a Spectronaut report based on the SPIN.rs scheme for DIA or extracted from the Maxquant output “evidence.txt” for DDA. Additionally, the aligned protein sequence database, the contaminant database, the experimental annotations, a list of species in the spectral library, and the fine grouping table are needed. The workflow is almost identical for the three types of data with small differences between MaxQuant and Spectronaut output column names, which can be found in the provided R scripts.

Species inference starts with loading the precursor identification files and filtering for 1 % FDR.

The databases are loaded and all isoleucines in the database and precursor sequences are changed into leucines. To allow for unambiguous site assignment, “global sites” are determined for every amino acid in the database by putting the 20 aligned genes in alphabetic order and numbering the positions from 1 through 25,550. The protein database is extended with an equal amount of “species decoy” proteins, which are generated by slicing the globally aligned sequences into 500 amino acid-long pieces and combining slices from randomized species into a new “chimeric” decoy species. The combined target/decoy database is used to annotate the precursors with matching genes, proteins, species, and global sites covered by the peptide sequence. Site-level information is generated by counting precursors/peptides, finding the maximum score, and summing the intensity for every possible amino acid at every possible global site, for each raw file. The summed intensities larger than 1 are log10-transformed and the precursor count, peptide count, log-intensity, and max. score are scaled by dividing by the maximum at the global site. These 4 metrics are then multiplied to calculate a joined score (J-score), which is again scaled by dividing by the maximum J-score at the global site.

Based on the combined target/decoy database, a site difference matrix is generated. For every possible comparison of 2 species, the difference matrix lists the global sites and amino acids that are different between them. Global sites with gaps in one of the species are ignored. For each raw file, these species-discriminating sites are then scored using the J-score. The species with the higher J-score sum is selected as the “winner”, while both species are “winners” in case of a “tie”. The best match for a raw file is the species that won most comparisons.

Fine grouping will be done for raw files with a best match that is amongst the species with manually selected marker peptides. The fine-grouping uses a list of hand-picked peptides that can be used to discriminate closely-related species within a genus. These peptides will be scored based on their precursor intensities in the respective sample. Based on the highest score, a single or multiple indistinguishable fine-grouping species will be reported.

The final output species is selected by taking either the best match or, whenever available, the fine-grouping species. The “site count”, which is the absolute sequence coverage, is added to this output table by counting the number of matching sites that were identified in the respective raw file. After ranking the list of raw files by decreasing site count, a q-value can be calculated as the fraction of decoy-species identifications at a given site count cutoff. Files with a q-value above 1 % will be marked in the final output. Furthermore, the relative protease intensity will be calculated for each sample by dividing the summed precursor intensity of proteases peptides by the total precursor intensity sum. The relative protease intensity cutoff is determined as the upper quartile of relative protease intensity amongst the laboratory blanks. Samples below that threshold will also be marked in the final output.

## Acknowledgements

We would like to express our gratitude to Daniel Klingberg Johansson and the Natural History Museum of Denmark for providing samples from non-human primates, Eske Willerslev for providing the mammoth bone, and Pedro Horta and Cláudia Costa for supporting the sample selection. We thank Alberto Taurozzi and Meaghan Emma Mackie for helping to explore the PAC methods for very ancient samples. Thank you to Ulises Hernández Guzmán for naming our method, and for Dorte Breinholdt Bekker-Jensen and Tanveer Singh-Baath for providing technical foundations for our workflow. P.L.R., E.C., and J.V.O. were supported by the European Commission through the MSC European Training Network ‘TEMPERA’ (grant number 722606). Work at The Novo Nordisk Foundation Center for Protein Research (CPR) is funded in part by a generous donation from the Novo Nordisk Foundation (NNF14CC0001). E.C. was supported by VILLUM FONDEN (no. 17649). F.W. has received funding from the European Research Council (ERC) under the European Union’s Horizon 2020 research and innovation programme (grant agreement No 948365) and a Marie Skłodowska Curie Individual Fellowship (No. 795569). The work in Vale Boi and Companheira is funded by Fundação para a Ciência e Tecnologia (FCT), grant PTDC/HAR-ARQ/27833/2017. The work at Lapa do Picareiro is funded by U.S. National Science Foundation (NSF) awards to J.H. (BCS-1420299, BCS-1724997) and M.B. (BCS-1420453, BCS-1725015). L.F. was supported by an SGS grant of the University of West Bohemia (SGS-2020-017). JC is funded by Fundação para a Ciência e para a Tecnologia (FCT), contract references DL57/2016/CP1361/CT0026. We thank all members of the JVO group for the constructive feedback, the MS platform team at CPR for keeping our instruments in good shape, and Thermo Fisher Scientific in Bremen for early access to MS instruments.

## Author contributions

P.L.R., J.V.O., and E.C. conceived the study. E.C. and F.W. established collaborations. P.B., K.M.G., P.P., M.C., R.M.G., L.F., J.C., M.L.S.J., M.M.B., J.H., N.B., F.W., and E.C. contributed archaeological samples. P.L.R., I.H., and F.W. performed laboratory research. P.R. and I.H. analyzed the data. P.B., K.M.G., P.P., M.C., R.M.G., L.F., J.C., M.M.B., J.H., N.B., F.W., and E.C. provided archaeological interpretations. P.R. prepared the manuscript. J.V.O., F.W., and E.C., and all co-authors contributed in the revision of the manuscript.

